# Bridge Category Models: Development of Bayesian Modelling Procedures to Account for Bridge Ordinal Ratings for Disease Staging

**DOI:** 10.1101/2021.08.17.456726

**Authors:** Joshua Levy, Carly Bobak, Nasim Azizgolshani, Michael Andersen, Arief Suriawinata, Xiaoying Liu, Mikhail Lisovsky, Bing Ren, Brock Christensen, Louis Vaickus, A. James O’Malley

## Abstract

Disease grading and staging is accomplished through the assignment of an ordinal rating. Bridge ratings occur when a rater assigns two adjacent categories. Most statistical methodology necessitates the use of a single ordinal category. Consequently, bridge ratings often go unreported in clinical research studies. We propose three methodologies (<u>Expanded, Mixture</u>, and <u>Collapsed</u>) *Bridge Category Models*, to account for bridge ratings. We perform simulations to examine the impact of our approaches on detecting treatment effects, and comment on a real-world scenario of staging liver biopsies. Results indicate that if bridge ratings are not accounted for, disease staging models may exhibit significant bias and precision loss. All models worked well when they corresponded to the data generating mechanism.

## 1 INTRODUCTION

Ordinal rating data are commonly used for routine clinical staging and grading of tissue biopsies[1, 2, 3]. However, raters may occasionally assign two adjacent categories, or bridge ratings. For instance, fibrosis in Non-Alcoholic Steatohepatitis (NASH), are staged by raters on a 5-point scale (from 0 to 4) [4, 5, 6] (Figure 1). While a stage 0 rating is likely to represent a healthy individual and stage 4 rating represent an individual with cirrhosis who may require liver transplantation, pathologists will sometimes assign bridge ratings (e.g. 2-3). Explanations for why such bridge ratings were assigned include the following motivating scenarios:

1. Expanded Scale (Figure 1B-C): Pathologist feels that an intermediate stage placed between the two adjacent stages would better encapsulate the disease pathology. However, since the intermediate category does not exist, they assign both categories.
2. Hedging by Blurring Stage (Figure 1D-E): Pathologist interprets scale incorrectly and rounds down or up erroneously on occasion. For instance, the pathologist may be told to report a stage 2 if they think the biopsy is a stage 2 with a probability of 0.7 and a stage 3 with a probability of 0.3, but may want to hedge against the potential of the more advanced stage being correct.
3. Collapsed Scale (Figure 1F): The pathologist believes the scale has one fewer category than available in the guideline scale, leading them to assign an interval that can compensate for information loss.

**FIGURE 1.**
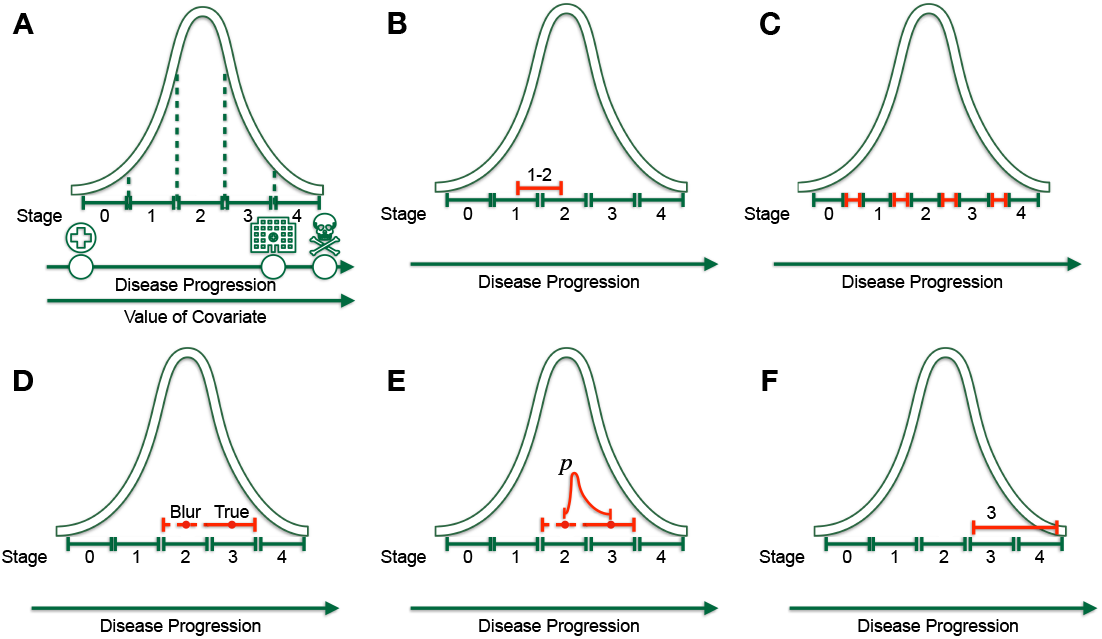
Visualization of data generating mechanisms and modeling approaches: A) Visualization of latent distribution/likelihood in cumulative link model, where top right arrow indicates latent scale of disease progression, while cutpoints/thresholds in latent distribution are denoted by vertical dashed lines. Areas between cutpoints correspond to rating/stage probabilities. Different predictors of the underlying latent rating may covary with latent disease progression, indicated by second right arrow. B) Data generating mechanism for expanded DGM, where biopsy may present features exactly between two adjacent ratings. The expanded DGM is accounted for using: C) the expanded category model, where additional categories indicated in red are estimated. D) Visual description of the blurred DGM, where here, true rating (*Y* = 3) is blurred into lower category (*Y* = 2 - 3). In the mixture adjacent model E) a mixture parameter p indicates the potential that the true rating is *Y* = 3. F) Visual description of the collapsed DGM, where the rater imagines scale is smaller than it really is and assigns a larger interval (*Y* = 3 - 4; red limits) based on the lack of information instead of *Y* = 3.

These explanations often arise in clinical research studies and practice. As common examples, sometimes the pathologist feels the tissue sample is sub-optimal in some way (e.g. too small, too fragmented, crushed, etc.), or the biopsy exhibits features of both stages. Alternatively, the pathologist may feels the clinical scenario does not match the histological findings (e.g. patient with sequalae of cirrhosis but liver biopsy showing only moderate fibrosis). In sum, bridge ratings may be employed to better describe the features of the biopsy.

In response to most statistical methodologies being unable to account for bridge ratings, domain experts often employ <u>ad hoc</u> approaches such as rounding the ratings to the higher or lower level, or randomly select either of the two ratings. Raters are often discouraged from reporting bridging categories and may resort to a combination of the aforementioned approaches [7, 8, 9, 10, 11, 12, 13]. In the practice of medicine, in addition to binary, categorical and continuous outcomes, the assessment of ordinal ratings is important for the conduct of clinical trials (e.g. screening, baseline and endpoint) [14, 15, 3], evaluation of the psychometric properties or other forms of validation of the underlying measurement scales (e.g. estimation of the intraclass correlation coefficient) [16, 5, 17], identification of important exposures/covariates [18, 19, 6, 20, 21], and validation of machine learning technologies which may predict an ordinal response [12, 22, 23, 24].

Some statistical methods exist for analyzing imprecise rating data under various assumptions about what the imprecise ratings mean[25, 26, 27, 28], but these methods are seldom employed in practice. We consider a situation in which the imprecise ordinal ratings could be interpreted in three different ways and we develop three statistical models appropriate for accounting for bridged ratings under each of these interpretations: 1) the expanded category model, 2) mixture adjacent model, and 3) collapsed category model.

We evaluate the performance of each of these statistical models under the different assumptions on the meaning of the bridge rating. This allows the evaluation of potential harm (e.g. bias, imprecision, high variance, erroneous coverage of interval estimators) from using the incumbent approach of rounding up, down, or randomly. We also assess which statistical model yields the most robust results in general. Finally, we present a real-world dataset (comparing serological markers and potential confounders to NASH fibrosis stage [12]) which contains bridge ratings and comment on the applicability of, and future directions for, our modeling approaches.

## 2 METHODS

In the appendix, we have included an introduction to ordinal regression models (e.g. ordered logit and ordered probit specifications) which are used to formulate our strategy for dealing with the bridged ratings of an ordinal response variable (section “Ordinal Response Variables and Cumulative Link Models (CLM)”). Let *Y* represent the ordinal response variable, *j* ∈ {1, 2,. . ., *K* - 1} (i.e., disease stage). Let *X* be a design matrix of observations by predictors. The latent variable, *Y*\*, represents the true, unobserved continuous process (i.e., disease progression) underlying the ordinal observations. The predictors serve to explain this progression/latent variable. With respect to disease staging, subjects with lower values in the latent scale are assumed to be healthier than those with higher values, who may exhibit significant progression. A pathologist who stages the disease may cut this continuous scale to form discrete stage measurements (Figure 1A). As per the derivation in the appendix, we implemented a fully Bayesian multinomial model using the Stan probabilistic programming language (See Appendix, section “Bayesian Computation and Hamiltonian Monte Carlo”) [29, 30, 31, 32] for disease staging. The cumulative link model (CLM) for *K* ordinal response categories is specified below for predictors indexed by *m* ∈ {1, 2,. . ., *M*} and cutpoints from *j* ∈ {1, 2, 3,. . ., *K* - 1}, under the strong assumption that effects (*β*) are invariant to *j* (not category-specific):

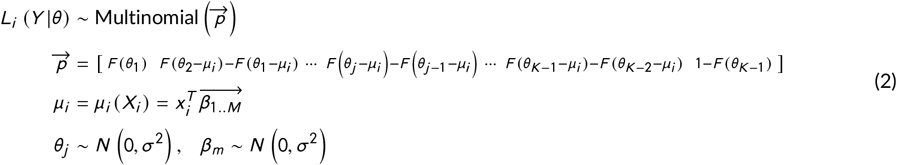

The following three subsections feature three separate data generating mechanisms (DGM) and corresponding statistical models, each of which builds upon the aforementioned model. In each of these subsections, we develop the likelihood function for each DGM and associated estimation procedure.

### 2.1 Description of Data Generating Mechanisms and Bridge Category Modeling Approaches

In response to the three motivating scenarios, we propose three adaptations of the cumulative link model to account for ratings of *j* and *j* + 1, which we denote as {*j, j* + 1}, in an ordinal rating scale with *K* categories. We denote the models developed herein for handling adjacent category data as ‘Bridge Category Models’, referring to the scenario where information pertaining to the assigned adjacent rating interval is censored (Figure 1). In all cases, we imagine an underlying latent distribution, *y*\* ~ *N* (0, 1) (under probit specification), is classified into *K* categories about *K* - 1 cutpoints.

### 2.2 The Expanded DGM: The Expanded Category Model

The <u>expanded category model</u> posits that the rater genuinely believes that the rating lies between <u>j</u> and <u>j+1</u> on the underlying continuous scale. This scenario may arise when the rater feels that the scale is too sparse and as such does not adequately distinguish between cases that the rater feels are genuinely different. As such, the rater reports a bridge rating as a way of conveying that the case is nestled between those typically represented by the bridged ratings. In the data generating model, ordinal data is generated through introduction of 2*K* - 2 cutpoints, where the area of the even integer categories in the latent space reflects the frequency at which uncertain assignments occur. We handle this by estimating a model with additional categories (Figure 1B-C). If such a scenario were to occur between all adjacent cutpoints when there are *K* possible ratings, the new scale would have 2*K* - 1 ratings, where the new categories bring added precision [33, 34, 35, 36, 37]. We assume here that all raters are thinking in terms of the 2*K* - 1 categories, which may differ from reality. Even numbered categories on this scale represent intervals on the original scale for which naming a single category is difficult. The expanded category model increases the resemblance to continuous data through introduction of additional categories which have smaller average distance between them than the original categories. It is common for ordinal response data with more than ten categories to be treated as continuous outcomes [33, 34, 35, 36, 37]. The specification of the Bayesian model is modified from the CLM model for *K* ordered categories for predictors indexed by *m* ∈ {1, 2, . . ., *M*} and cutpoints from *j* ∈ {1, 2, 3,. . ., 2*K* - 2}:

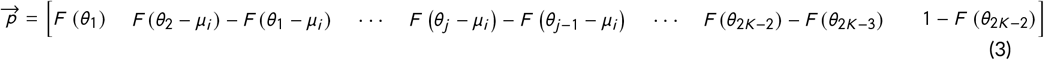

The priors and likelihood are of the same functional form as the original CLM. The distance between these intermediate cutpoints and the immediately adjacent cutpoints indicates the relative frequency with which bridge-ratings were made by raters.

### 2.3 The Blurred DGM: The Mixture Adjacent Model

Here, we consider the case when two adjacent ratings are blurred. Data generated from the latent distribution is organized into *K* categories via *K* - 1 cutpoints. A proportion of the ratings, *q*, are blurred, where for each rating *j*, the rating reported is {*j, j* + 1} with probability p and {*j* - 1, *j*} with probability 1 - *p*. This corresponds to the true rating being the smaller rating of the pair *p* proportion of times and the bigger rating 1 - *p* proportion of times.

The <u>mixture adjacent model</u> posits that the rating was mis-coded, or there were defining features in either the higher or lower rating that made a combined rating more appealing (e.g. most of the histological features resemble a stage two, but some are indicative of a stage three; “mostly stage two”). In this scenario, the true solution could be either <u>j</u> or <u>j+1</u>, but we are unsure of which. We denote the probability that the true rating is the lower rating as <u>p</u>. The likelihood function for the entire sample of observations is given by:

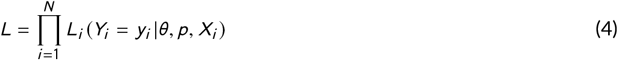

with the contribution from the ith observation given by:

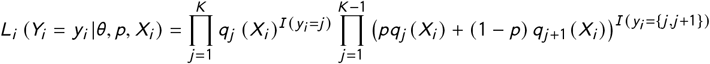

where,

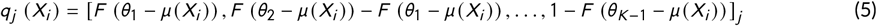

with F() the cumulative distribution function of the standard normal distribution,

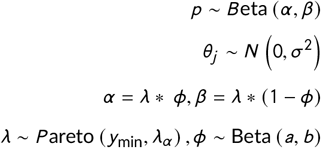

A prior in the beta family is set on the mixture parameter, *p*, and accommodates prior knowledge on whether the population behavior favors leaning towards down or up-rating. Hyperpriors *λ* and *ϕ* are mean and count parameters that may be reparameterized as the parameters of the beta prior (*α*, *β*). Alternatively, if the mixture parameter is assumed apriori, we set *p* to be some constant between 0 and 1, which we refer to as the <u>Set Mixture Model</u>. This DGM could arise in practice if the measurement system had a known property that led to an adjacent rating erroneously appearing to be the correct rating a certain proportion of the time (e.g., system fault). However, it is not as flexible as one which does not impose constraints on the mixing probability: it requires perfect knowledge of the mixing probability. The mixture parameter, proportion <u>p</u>, should be recoverable by the <u>Mixture Adjacent Model</u>.

### 2.4 Collapsed DGM: The Collapsed Category Model

While the <u>Mixture Adjacent Model</u> handles blurred measurement, we expect the <u>Collapsed Category Model</u> to have some flexibility to the blurred categories. The <u>collapsed category model</u> refers to a data generating mechanism where the scale contains levels that the rater is unable to distinguish. As an example, the rater may report *Y* = {*j, j* + 1} as a single category as they are unable to distinguish between the categories. Analogous to interval censoring, we handle this scenario by combining the adjacent categories in the contribution from this particular rater to the likelihood function:

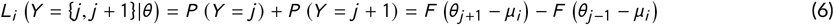

The likelihood of the Bayesian multinomial model (eq. (2)) may by modified to encapsulate this observation:

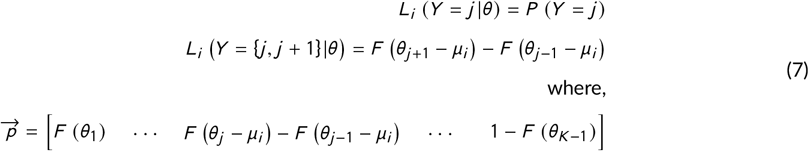

### 2.5 Description of Simulation Studies

We designed simulation studies to evaluate the benefit of correctly accounting for bridge-ratings and the robustness of the above three models to erroneously assuming the meaning of bridge ratings. We focus on the recovery of parameters of interest given a causal model. In all simulations, we evaluate the performance of the point and interval estimators of the effect of three covariates *X* [38], as they pertain to generation of latent information *Y*\*, which is turned into five-to-nine ratings categories (depending on the model generating the data) by thresholding the generated latent distribution (Figure 2).

**FIGURE 2.**
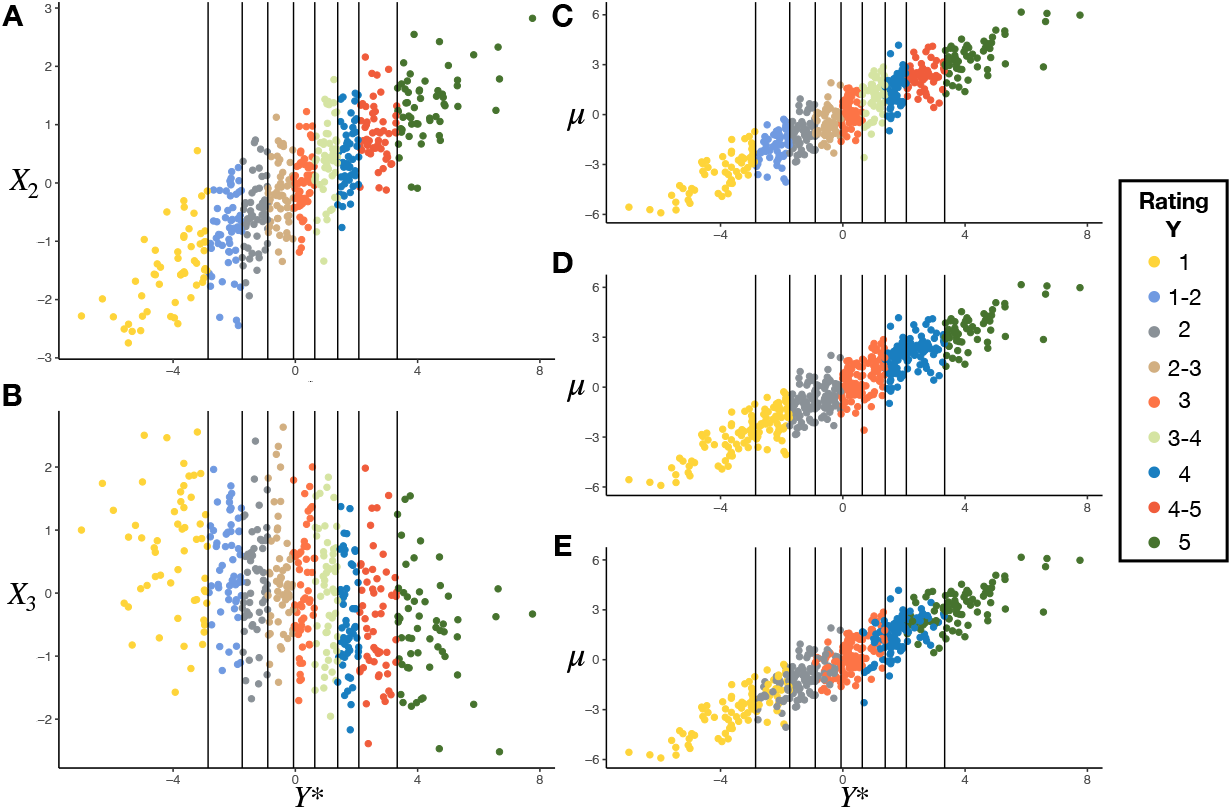
Demonstration of data simulation for the expanded DGM under various statistical / ad-hoc modeling procedures: vertical lines represent cuts in the latent distribution and points are colored by assigned ordinal rating: A) latent distribution *y*\* versus *X*_2_; B) latent distribution *y* * versus *X*_3_; C) latent distribution *y*\* versus conditional mean *μ*; D) example of down-rating uncertainty categories (even numbered); E) example of random-rating uncertainty categories; rating Y=1 indicates early/healthy staging for individual, while categories closer to Y=5 indicate later/severe staging

### 2.6 Data Generation: True model Known and Referred to as Data Generating Model (DGM)

The first covariate was simulated from a uniform distribution and thresholded to form a binary covariate with a prevalence of 0.2 while the remaining covariates were simulated from a standard normal distribution such that *Z* ~ *U* (0, 1), 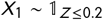, *X*_2_ ~ *N* (0, 1), *X*_3_ ~ *N* (0, 1). These covariates correspond to scenarios where one may be estimating a treatment effect (binary covariate) of a marker of interest or wanting to adjust for a continuous confounder (continuous covariate). Covariate effects are given by:

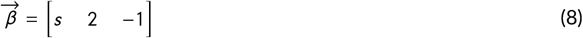

Where *s* is a sensitivity parameter (true effect size of binary covariate) under various simulations while the other parameters remain fixed. Given a probit link function, data is simulated as:

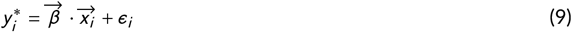

where *ϵ*_*i*_ ~ *N* (0, 1).

Here, cutpoints 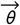 are established to cut this distribution into *n* ordinal ratings, *Y*. Then, observations are augmented to conform to a specific DGM. To generate bridge ratings under the expanded DGM, 2*K* - 2 thresholds are utilized for the binning procedure. The probability mass of each of the even integer categories (uncertain categories; e.g. 2-3) is denoted by *e*, which can be tweaked from 0 (no uncertainty) to 0.25 (100% uncertainty). To generate bridge ratings under the blurred DGM, ratings are blurred into adjacent categories. We denote the amount of blurriness (proportion of times original assignment is blurred into uncertain adjacent categories), of the Blurred DGM as *q*. Conditional on having blurred observations, the proportion of times in which the blur is of the higher adjacent categories is denoted as *p*. Cutpoints are chosen such that equal probability is assigned to each rating prior to application of the blur mechanism.

We did not generate data under the collapsed DGM. Based on interviews with domain experts, the scenarios by which data may be generated in a measurement scale with a lower number of ratings and inferred to be in a scale with a greater number of categories may be less applicable and intuitive across biomedical research domains.

### 2.7 Model Estimation: True Model (or DGM) Unknown

For each simulation, after binning observations into ordinal and bridge ratings, we estimate the main effects and cutpoints for each DGM (expanded, blur) under the following data augmentation and modeling procedures:

1. Up-Rating: All “blurred” or “adjacent” assignments produced from the expanded or blurred DGM are assigned to the higher of the two ratings. An ordinal regression model with *K* ordinal ratings is fit.
2. Down-Rating: All “blurred” or “adjacent” assignments produced from the expanded or blurred DGM are assigned to the lower of the two ratings. An ordinal regression model with *K* ordinal ratings is fit.
3. Random-Rating: All “blurred” or “adjacent” assignments produced from the expanded or blurred DGM are randomly assigned to either the higher or lower of the two ratings. An ordinal regression model with *K* ordinal ratings is fit.
4. Expanded: Fitting the Expanded model under the assumption of 2 × *K* - 1 categories.
5. Mixture Adjacent: Fitting the Mixture Adjacent Model.
6. Collapsed: Fitting the Collapsed Model.

We term the first three approaches as the <u>traditional approaches</u>, since they represent how bridged ordinal ratings had previously been handled. The Up-, Down-, and Random-rating scenarios (analysis methods 1-3) correspond to the true DGM under extreme special cases of the blurred DGM (*p* = 0, *p* = 1, and *p* = 0.5, respectively). These provide comparative performance criteria for the models in 4-6 and especially to the mixture adjacent model in 5, which encompasses them. To demonstrate differences between the traditional and bridge category modeling approaches, we consider the following simulation-based evaluation of the performance of each modeling approach under each specified DGM, fixing the number of categories to *K* = 5:

1. *e* ∈ {0, 0.05, 0.1, 0.15, 0.2} (Expanded DGM; *s* = 1)
2. *e* ∈ {0, 0.05, 0.1, 0.15, 0.2} (Expanded DGM; *s* = 6)
3. *n* ∈ {100, 200, 500, 1000, 2000} (Expanded DGM)
4. *s* ∈ {1, 3, 5, 6} (Expanded DGM)
5. *q* ∈ {0, 0.3, 0.5, 0.8} (Blurred DGM)
6. *p* ∈ {0, 0.3, 0.5, 0.8, 1} (Blurred DGM)
7. *n* ∈ {100, 200, 500, 1000, 2000} (Blurred DGM)
8. *s* ∈ {1, 3, 5, 6} (Blurred DGM)

The default parameters (the values assumed unless stated otherwise) are 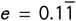, *q* = 0.6, *p* = 0.5, *n* = 1000, *s* = 1 (for blurred DGM), and *s* = 6 (for expanded DGM, where proportions of binary covariate in adjacent categories may vary greatly). When *e* =0 or *q* = 0, this is equivalent to the scenario from which no uncertainty or measurement error is introduced into the DGM.

For each sensitivity analysis, we generated 100 different datasets and estimated the posterior distribution of the covariate and mixture parameters (when using the <u>Mixture Adjacent Model</u>). We recorded posterior means and 95% high density credible intervals for these four parameters.

From these estimates, we provide frequentist estimates of the performance of these modeling approaches through reports of the bias and mean squared error (MSE) of the posterior mean compared to the true parameter values, coverage (percentage of times where credible interval covers the true population parameter, which should be close to the nominal 95% probability) and averaged posterior width (the utility of the interval estimator). Simulation analyses were performed on the Discovery Research Computing cluster at Dartmouth College.

### 2.8 Selection of Prior Distributions for Simulation Studies

We set the default values for the aforementioned priors and hyperpriors for all simulations to be *σ*^2^ = 1000, *a* = 1*e*^8^, *b* = 1*e*^8^, *y*_min_ = 0.1, *λ*_*α*_ = 1.5.

Given the constraints on the priors and hyperpriors, the priors over the cutpoints and covariate parameters were uninformative (*σ*^2^ is high). Meanwhile, the prior over the mixture parameter was centered around 0.5, given heavy centering of the hyperprior over the mean hyperparameter, The spread of the hyperprior was commensurate with the specified pareto distribution [39]. Decreasing *a, b*, and *λ*_*α*_ favors mixture parameter sampling towards the tails of the beta distribution. Increasing *y*_min_ may do the same but truncates the Pareto hyperprior and can introduce divergent geometry into the posterior.

### 2.9 Description of Real-World Dataset and Final Experimental Design

To test the external applicability of such models, we acquired a dataset of 286 steatohepatitis liver biopsies staged for fibrosis (featured in a previous validation study [12]) with fibrosis staging from four independent raters with a re-test and staging of an alternative modality for presenting liver biopsy images (a total of three fibrosis measurements per rater, 24 measurements per biopsy). Some subjects had multiple biopsies. We excluded the ratings from the alternative modality (16 measurements per biopsy), and simplified the scenario by considering information from one randomly selected biopsy from each subject. We selected three out of the four raters that had the lowest test-retest reliability and for each rater, selected the test which had the highest degree of adjacent assignments and fit all models (1-6) for each rater and compared the fits within rater. We assessed the pathologists separately to both match the low complexity offered by the simulation studies and understand how the six different modeling approaches produce different effect estimates within each rater.

Serological markers known to correlate with fibrosis staging also were measured in subjects with liver biopsy [40, 12, 41, 42]. We modelled fibrosis stage as the ordinal response, regressing on the Fib4 score (an estimate of the amount of scarring in the liver; 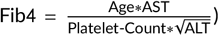) and the AST:ALT ratio (a proxy for the degree of liver dysfunction through two markers of hepatocellular injury), adjusting for BMI as a potential confounder. Predictors were centered and divided by their standard deviation prior to fitting each model in order to compare each predictor’s relative importance and assist with sampler convergence. We fit each model given the six data augmentation and modeling strategies featured above for the three covariates. We expected mean effect estimates for the three covariates to vary between raters due to interrater variability, though we leave exploration of the impact of bridge category modeling on rater intercepts to follow-up work.

### 2.10 Software Availability and Alternative Modeling Approaches

Simulation code (available in R 3.6 [43]), data preparation, Stan models, and additional scripts to assist with fitting real world data are available on GitHub at: https://github.com/jlevy44/BridgeCategoryStagingModels. Regression models are of the multinomial family, under the ordered probit (featured in this work) and logit specifications. Two alternative Stan fitting procedures correspondent to the <u>Mixture Adjacent Model</u> are provided in the simulation code which may be helpful for debugging as they are special cases of the general model. The former sets *p*, the mixture parameter, and does not seek its estimation. The latter estimates *p* by first drawing from a Bernoulli distribution parameterized by probability *p*, then using the draw to indicate whether to evaluate the likelihood of the lower or higher category. Averaging across all posterior draws should yield similar estimates to the <u>Mixture Adjacent Model</u>.

## 3 RESULTS

### 3.1 Simulation Studies

To evaluate where each approach may be of greatest benefit, we conducted approximately 30,000 simulations (100 simulations per configuration; approximately 900 configurations of sensitivity parameters, data generating mechanisms, modeling approaches). Estimates of the binary covariates are of particular interest and are shown in the upper left panel of the figure boxplots, which display the distributions of mean posterior estimates across the simulations for each modeling approach. We have included a tabular breakdown of the effect estimates in the appendix and an additional file (Appendix/Additional File 1; “Summary”).

### 3.2 Simulations for which the Expanded DGM is the True Model

Summarized reports of effect estimates and their agreement with the ground truth for each modeling approach fit after varying the degree of uncertainty for small (Table “Expanded-Degree-Uncertain(S=1)”) and large effect sizes (Table “Expanded-Degree-Uncertain(S=6)”), the sample size (Table “Expanded-Number-Samples(S=6)”) and size of the true effect estimate (Table “Expanded-Effect-Size”) of the binary covariate can be found in Appendix/Additional File 1 (“Summary”).

#### 3.2.1 Effect of Degree of Uncertainty on Model Performance when Expanded DGM is the True Model

When none of the assignments were denoted as uncertain, the expanded, up, down, and random staging approaches all yielded similar performance with low bias, low MSE and high coverage of the true effects (Figure 3, Appendix Figure 1). As we increase the amount of uncertainty to where the number of ordinal assignments were similarly distributed between certain and uncertain categories, the expanded, up, down, and random staging approaches yielded similar bias estimates (Figure 3, Appendix Figure 1). However, as expected from the theory of efficient estimation the interval width of the credible interval and the MSE are substantially lower for the expanded approach, the correct model, versus the traditional approaches. The difference between the expanded and traditional approaches are substantially greater for larger effect sizes of the binary covariate (Figure 4, Additional File 1). At higher effect sizes and uncertainty of *e* = 0.1, coverage for the binary covariate is greater for the expanded versus the traditional approaches. The MSE for the traditional approaches continues to climb given more uncertainty in ordinal assignments (Figure 4). While bias for the up, down, and expanded approach is negligible across uncertain assignments, bias for the random, mixture and collapsed approaches increases in magnitude substantially and coverage decreases. The mixture model was able to recover the true effect estimates at the greatest degree of uncertainty (*e* = 0.2).

**FIGURE 3.**
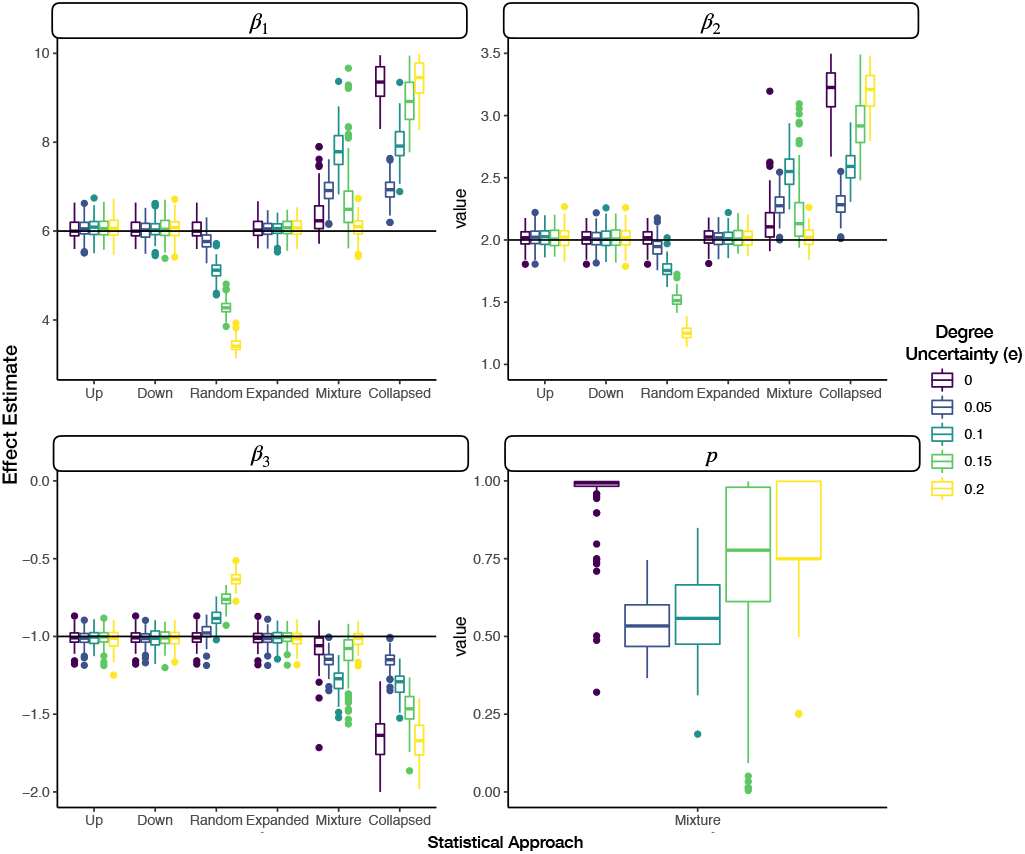
Faceted grouped boxplots indicating distribution of mean posterior covariate and mixture parameter effect estimates across simulated datasets; each facet presents different covariate/mixture effect; x-axis labeled by which data augmentation/modeling approach was fit to the data; y-axis is effect estimate; horizontal line added to indicate true population parameter for each covariate; boxplots are colored by the degree of uncertainty in the expanded DGM, where the effect size of the binary covariate is 6

**FIGURE 4.**
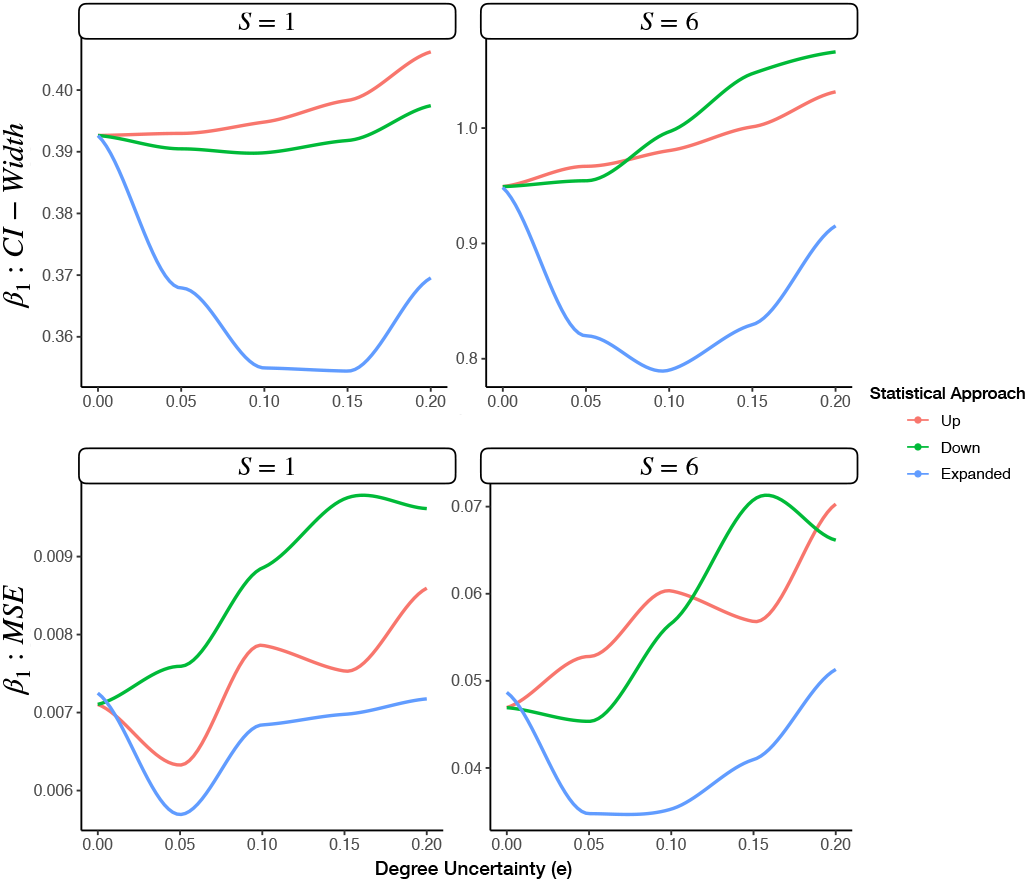
Degree of uncertainty (e) versus posterior interval width and MSE for the up, down and expanded category models for a true effect size of one for the left two plots and a true effect size of six for the right two plots; results reported for first covariate and smoothed using loess regression to better portray relationships

#### 3.2.2 Effect of Sample Size on Model Performance when Expanded DGM is the True Model

At low sample sizes, the expanded category model provides more precise estimates than the traditional approaches in terms of the average posterior interval width and lower MSE (Appendix Figure 2–3). The downstaging approach experiences high separation of the binary covariate between adjacent categories for some of the simulations at a low sample size, resulting in a high effect estimate that is highly biased. At higher sample sizes, the differences in precision and MSE between the expanded approach and its up/down counterparts are less apparent (Appendix Figure 3). Meanwhile, the bias of the random, mixture and collapsed approaches appears to approach different fixed values at these higher sample sizes (Appendix Figure 2).

#### 3.2.3 Increasing the True Effect of Binary Covariate on Model Performance when Expanded DGM is the True Model

Across all effect sizes of the binary covariate for the expanded DGM, the expanded model demonstrates lower posterior interval width and MSE versus down and up staging. As the effect size increases, so does the difference in interval width and MSE. Increases in interval width across effect sizes can be attributed to the larger magnitude of effects being measured, of which absolute deviation from the true value may increase, and potentially more sensitive imbalance of the binary covariate across the ordered categories. At the largest effect size, there is around a 20-30% reduction in interval width and MSE by the expanded as compared to up/down rating (Figure 5, Appendix Figure 4).

**FIGURE 5.**
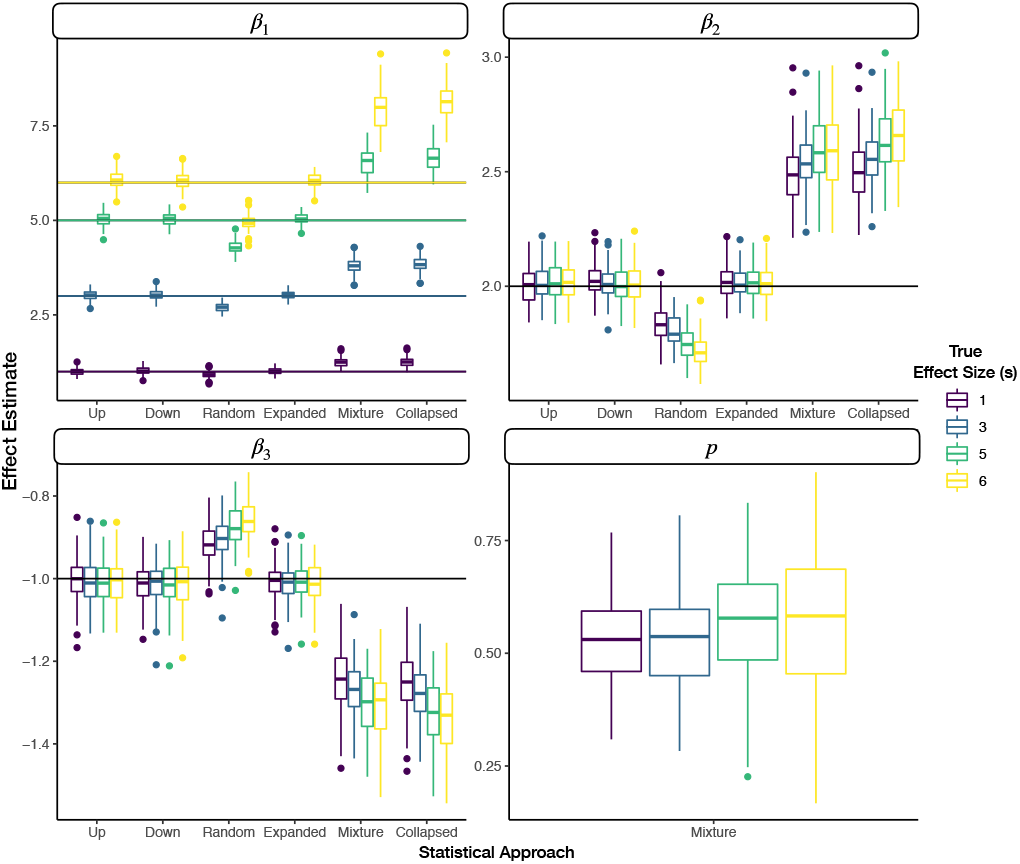
Faceted grouped boxplots indicating distribution of mean posterior covariate and mixture parameter effect estimates across simulated datasets; each facet presents different covariate/mixture effect; x-axis labeled by which data augmentation/modeling approach was fit to the data; y-axis is effect estimate; horizontal line added to indicate true population parameter for each covariate; boxplots are colored by the true effect size (s) of the binary parameter (*β*_1_) the expanded DGM, where the color of the horizontal line in the first panel indicates the true effect size

### 3.3 Simulations for which the Blurred DGM is the True Model

We now consider the case when the mixture model corresponds to the true DGM and assess its performance across various settings of the data and model parameters as well as the robustness of the other models, which are incongruent with the DGM. Summarized reports of effect estimates and their agreement with the ground truth for each modeling approach fit after varying the degree of blurring (Table “Blur-Degree-Up(S=1)”), degree of up-rating for blurred categories (Table “Blur-Degree-Up(S=1)”), the sample size (Table “Blur-Sample-Size(S=1)”) and size of the true effect estimate (Table “Blur-Effect-Size”) of the binary covariate can be found in the Appendix / Additional File 1.

#### 3.3.1 Effect of Degree of Blurring on Model Performance when Blurred DGM is the True Model

With increases in the degree of blurring introduced to the ordinal outcomes, we obtain higher bias (degradation of the effect magnitude) and MSE estimates for the expanded, up, down and random staging models, with substantial reductions in coverage. The magnitude of the effect for these approaches begins to decrease towards zero given this blurring. In contrast, the collapsed category and mixture models are able to recover the true effect with optimal coverage (around 0.95, the nominal probability) for all degrees of nonzero blurriness (Figure 6, Additional File 1). We found negligible bias and MSE for these models versus the other approaches. However, the average width of the 95% credible interval is higher for these two models versus the other modeling approaches, which may be an artifact of providing larger magnitude effect estimates and how the covariate is distributed across the categories as the effect becomes larger. The mixture model is also able to recover the true mixture population parameter with high coverage (~0.91 coverage).

**FIGURE 6.**
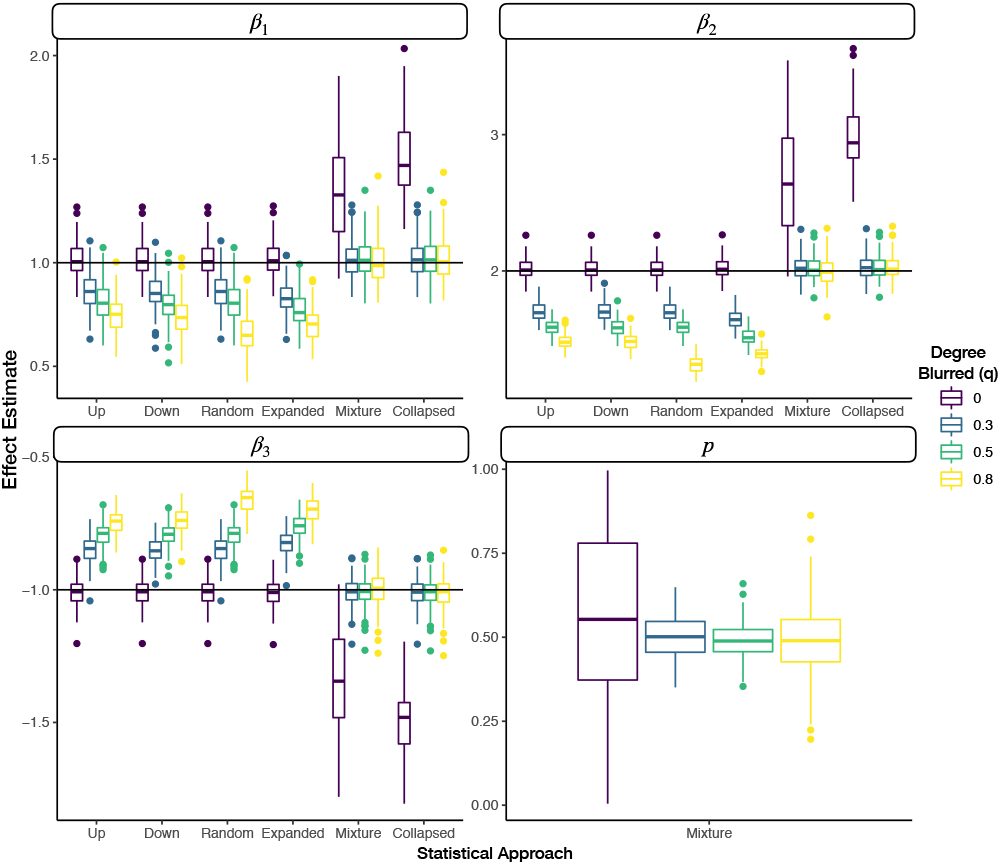
Faceted grouped boxplots indicating distribution of mean posterior covariate and mixture parameter effect estimates across simulated datasets; each facet presents different covariate/mixture effect; x-axis labeled by which data augmentation/modeling approach was fit to the data; y-axis is effect estimate; horizontal line added to indicate true population parameter for each covariate; boxplots are colored by the degree of blurring (q) in the blurred DGM

#### 3.3.2 Effect of Proportion of Higher Assignments on Model Performance when Blurred DGM is the True Model

When varying the proportion of assignments that were blurred to the upper two adjacent categories (<u>p</u>), the expanded and random rating models demonstrate high bias, high MSE, low coverage estimates, which drifts towards a zero-magnitude effect estimate. For the up-staging models, performance is optimal when *p* = 0 (high coverage, low bias and MSE, slightly higher precision than the mixture approach). As *p* increases, the bias (away from the true parameter) and MSE increase and coverage quickly drops. For the down-staging, the opposite holds true: optimal performance / recovery of the true effect when *p* = 1; increases in bias (away from the true parameter) and MSE occur while there is a decrease in coverage as *p* decreased.

The mixture model recovers the true covariate effect at all values of *p*, with high coverage, low bias and MSE (Figure 7, Additional File 1). In all cases, the mixture model is also able to recover the true mixture population parameter with high coverage close to the nominal 95% probability, low bias and MSE. This holds true except for the cases where *p* = 0 or *p* = 1, where coverage is 0. However, this is an artifact of the impossibility of the posterior mean of the parameter to be equal to precisely 0 or 1, under a prior distribution that is a continuous density which places 0 support on both *p* =0 and *p* = 1. The reductions in posterior interval, bias and MSE near the tails suggest that the true parameter has effectively been recovered. The collapsed model exhibits performance similar to that of the mixture model. However, performance is greatest when *p* = 0.5, where there is no predisposition towards up/down-staging. Coverage remains high for the binary covariate but is slightly reduced when *p* approached 0 or 1. The magnitude of the bias and MSE also increases slightly as *p* approached 0 or 1 (Figure 7, Additional File 1).

**FIGURE 7.**
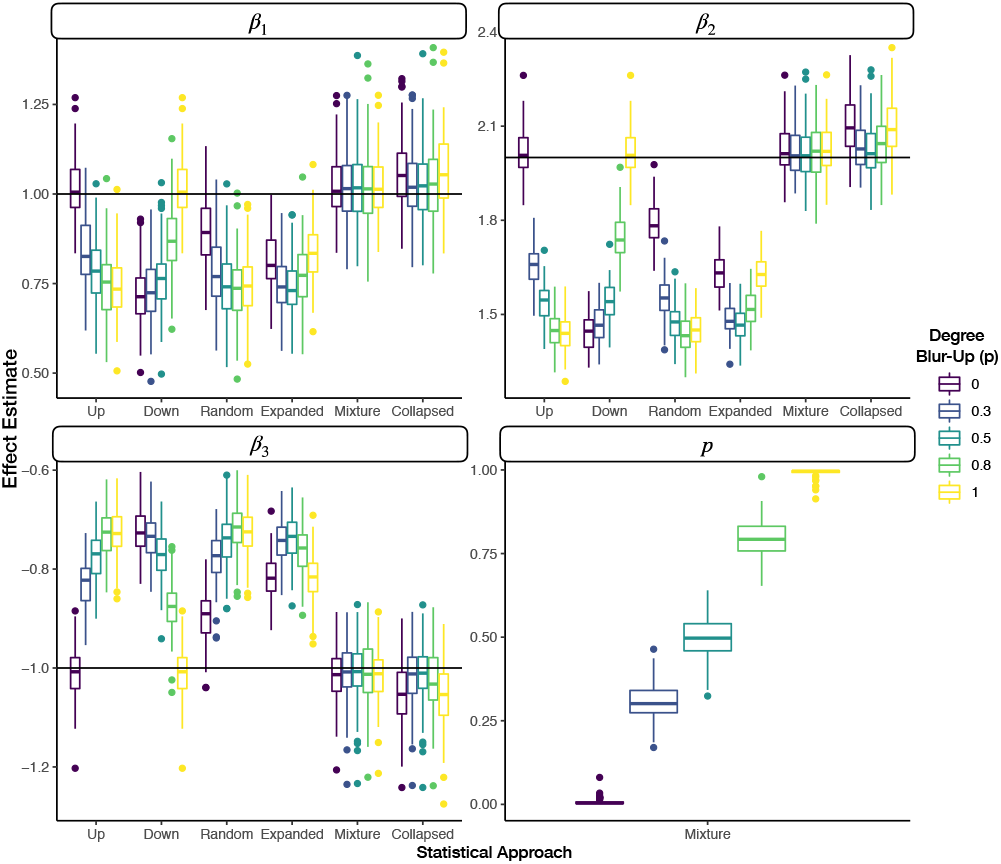
Faceted grouped boxplots indicating distribution of mean posterior covariate and mixture parameter effect estimates across simulated datasets; each facet presents different covariate/mixture effect; x-axis labeled by which data augmentation/modeling approach was fit to the data; y-axis is effect estimate; horizontal line added to indicate true population parameter for each covariate; boxplots are colored by the proportion of times when the true rating was blurred into the upper two ratings (p) via the blurred DGM

#### 3.3.3 Effect of Sample Size on Model Performance when Blurred DGM is the True Model

Increasing sample size is associated with decreases in posterior width for the covariate and mixture parameters across all models. For the expanded, down, up and random models, the reductions in posterior width is correspondent to reductions in coverage given the already biased estimates of the approaches. Meanwhile, coverage for the mixture and collapsed models remain largely unaffected. Finally, the mixture parameter is recovered with high coverage and decreasing posterior credible interval width for higher sample sizes (Appendix Figure 5, Additional File 1).

#### 3.3.4 Increasing the True Effect of Binary Covariate under the Blurred DGM

The mixture and collapsed category models recover the true effect across the full range of effect sizes (Figure 8, Additional File 1). The magnitude of the effect estimates under the remaining approaches are significantly below their target value across all effect sizes of the binary covariate. Coverage of the expanded, random, up and down stage models is close to 0 while there is nearly nominal coverage (close to 95%) of the covariate and mixture effects using the mixture and collapsed model approaches.

**FIGURE 8.**
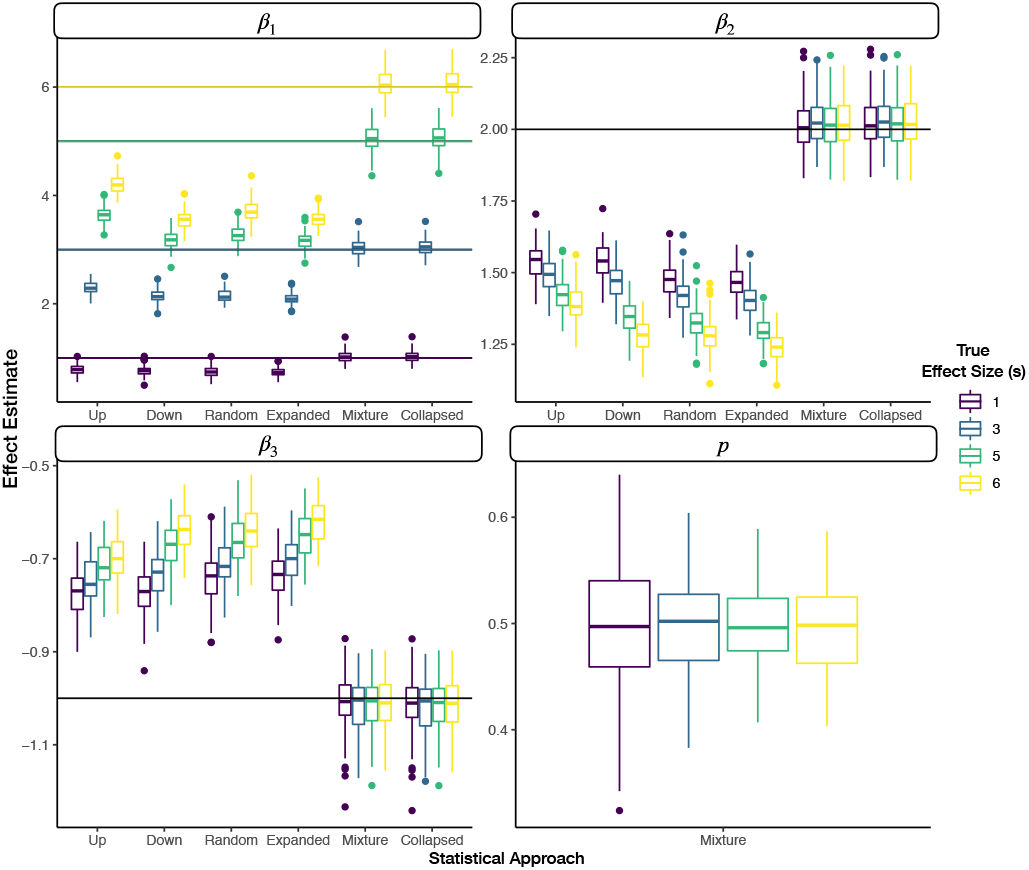
Faceted grouped boxplots indicating distribution of mean posterior covariate and mixture parameter effect estimates across simulated datasets; each facet presents different covariate/mixture effect; x-axis labeled by which data augmentation/modeling approach was fit to the data; y-axis is effect estimate; horizontal line added to indicate true population parameter for each covariate; boxplots are colored by the true effect size (s) of the binary parameter (*β*_1_) under the expanded DGM, where the color of the horizontal line in the first panel indicates the true effect size

### 3.4 Real World Application

We now report the effect estimates and 90% highest posterior density credible intervals for various standardized measurements versus the assigned fibrosis stage (Table 1, Appendix Figure 6, Appendix Tables 1–2). Between the two pathologists, the proportion of two-rating adjacent stage assignments over all assignments are 0.30 and 0.25 for pathologists 1 and 2 respectively. Between the raters, the AST:ALT ratio is given the highest association with Fibrosis progression, followed by the Fib4 score (based on pathologist 2; Appendix Figure 6B), then BMI. These effects vary significantly depending on which rater was assigning stages and which model is estimated. Between raters and models, the mean posterior estimate of BMI varies from 0.107 to 0.164. For Fib4, mean estimates range from 0.0702 to 0.195. For the AST:ALT Ratio, mean effect estimates range from 0.2 to 0.414.

**TABLE 1.**
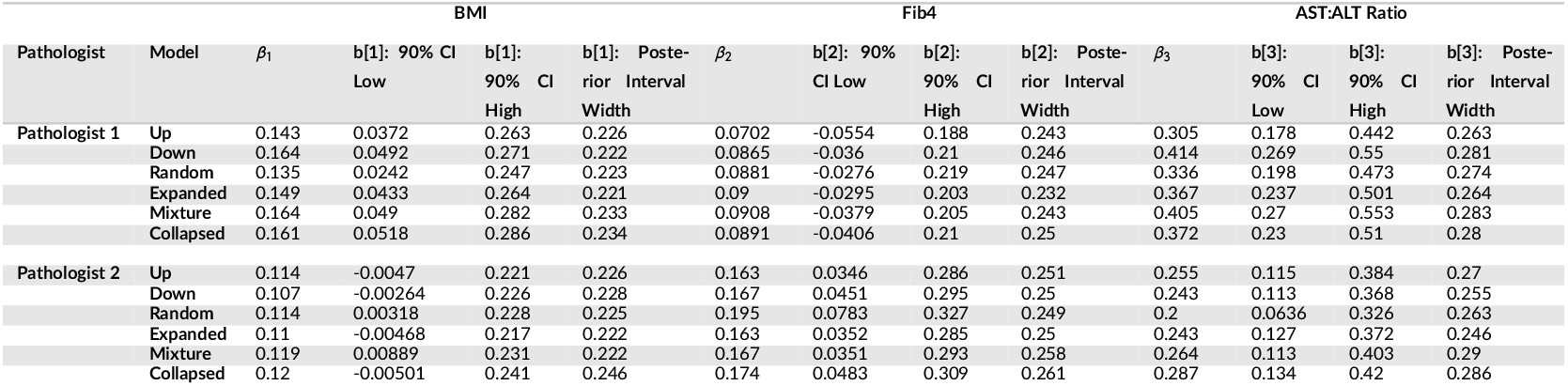
Posterior estimates for covariates for fibrosis staging model (BMI [1], Fib4 [2], AST:ALT Ratio [3])

Under the assumption of the blurred DGM as the true DGM, pathologist 1 (mixture parameter 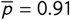) demonstrates preference for assigning the higher adjacent stages while the lower stage was true. In comparison, pathologist 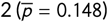 prefers assigning the lower adjacent stages while assuming the upper stage was true. If the Blurred DGM was the truth, covariate effects determined by the mixture model are similar to the down-staged model for pathologist 1 and similar to the up-staged model for pathologist 2. For these raters, these effect estimates appear to differ from upstaging. For pathologist 2, the effect of BMI on Fibrosis becomes positively significant according to a 90% credible interval for the mixture model.

## 4 DISCUSSION

Clinical grading and staging scales, largely ordinal in nature, are incredibly important for the assessment of disease, not only for real-time clinical decision making, but also for establishing screening and assessment of baseline predictors and endpoint outcomes for the conduct of FDA regulated drug trials. The existence of measurement error, uncertainty and reliability issues between raters may reduce the study power, thereby causing a drug to fail clinical trials [44, 45]. While many rating scales, such as the NASH CRN scale for the grading and staging of liver fibrosis for progression into cirrhosis, have been validated through testing of interrater reliability [17], it is not uncommon to see follow-up studies dispute the reported reliability [15, 5]. Consequently, the effectiveness of the scale itself may be called into question as it pertains to these aforementioned matters (clinical triage and trials).

The assignment of two adjacent ordinal ratings is no different. Although under-reported, the potential impact of such ratings on interrater reliability and measurement of a treatment effect across the appropriate biomedical discipline should be explored within other biomedical specializations. For instance, the AJCC Cancer Staging Manual recommends for TNM (tumor histology, lymph node, metastasis) staging that in the presence of uncertain assignment of stage, the lower of two possible adjacent categories should be assigned [7, 10], while other interpretations of the guide have pointed out that the highest stage descriptor should be selected [46]. For staging criteria with subcategorization (e.g. Stage 1a, 1b, 1c), the AJCC guide also recommends assigning the general category (e.g. Stage 1) and reporting that the tissue cannot be assessed[7]. In this scenario, the rater would cast uncertainty over the three ordinal measurements contained within the staging group (e.g. Stage 1). These guidelines have informed staging practices internationally, yet some critics have pointed out that these recommendations regarding stage uncertainty are not substantiated and leave room for mis-classification [13]. Furthermore, we suspect that active discouragement for reporting ambiguity in staging can lead to the application of these <u>ad hoc</u> procedures or potentially the omission of the observations and may explain why such reports are virtually ignored in the research setting.

In this paper, we discussed the treatment of simultaneous assignment of two adjacent ordinal ratings, bridge ratings, and potential implications when evaluating the effect of binary (eg. treatment) and continuous (eg. some serological marker or confounder) covariates. Clinicians indicate in interviews that simultaneous adjacent stage assignments are a common occurrence, yet they are often obscured from the view of the reader/regulator because methods to account for these uncertain adjacent assignments do not currently exist. We developed three likelihood functions (the expanded, mixture adjacent, and collapsed bridge category models) and evaluated for their potential to yield meaningful results when the expanded and blurred DGM are the true models. We ran simulation studies to illustrate where to expect the greatest advantage when using the expanded, mixture and collapsed approaches versus the traditional approaches under the expanded and blurred DGM. We followed this analysis with a real-world use case, identification of clinical and serological factors pertaining to the staging of liver fibrosis for the assessment of NASH.

From our simulation studies of the expanded and blurred DGM, we found that the bridge category models were able to outperform the traditional approaches with respect to their applicable data generating mechanism.

The results from the simulation studies highlight that it is critical to select a modeling approach commensurate with the true data generating mechanism. Misalignment of the approach with the DGM can result in more biased and/or less precise estimates. For instance, we noticed that the mixture and collapsed models were not well-adjusted to the expanded DGM and biased effect estimates with reports of higher magnitude effects than intended. Meanwhile, the expanded DGM suffered the same deficiencies that the up/down/random staging models experienced on the blurred DGM. We note that up and down rating were relevant for particular use cases (*p* = 0 and *p* = 1 respectively). These may be special cases where the mixture model can essentially learn whether the augmentation applied to the data should be up or down-staged. The effects of the misalignment are far reaching in the small sample setting, while coverage can rapidly diminish in the large sample setting with improper treatment of the bridge rating.

From our real-world models fit to estimate fibrosis stage under the probit specification, it was difficult to characterize the true DGM. However, we noticed a tendency of effect estimates to track that expected of lower and higher staging, as suggested by the effects from the mixture modeling approach. This information is concordant with interviews conducted with pathologists at the Department of Pathology at Dartmouth Hitchcock Medical Center, where pathologists were asked for reasons that could explain the bridged assignments. One pathologist indicated the biopsy had features which were mostly indicative of the lower of two stages, while some features suggestive of the higher of the two adjacent stages were also present: “When I use [bridge ratings], i.e. stage 2-3, the specimen shows features that are mostly indicative of a stage 2, though focal area with possible bridging fibrosis are suggestive of a stage 3. The specimen is not a definitive stage 3. The same is true for a stage 3-4, where the specimen shows features that are predominantly stage 3 (bridging fibrosis), though focal areas will show nodule formation that are suggestive of stage 4. The specimen is not a definite stage 4”. We received feedback from another pathologist, who had noted that the NASH CRN scale had been developed with an abundance of tissue material with multiple portal regions from which to make stage assignments [47, 5, 48]. For clinical trials, biopsies which may contain ample portal tract (at least 10 tracts), needle biopsies, and large tissue cores (2-3 cm) are recommended[48]. The pathologist suggested that in practice, liver biopsies often contain scant information. As such, an incomplete set of diagnostic features may diminish the confidence in stage assessment. Finally, outside of the host institution, another pathologist had indicated to our group that grading and staging assessment is non-uniform across the tissue biopsy [49, 50, 51]. In summary, a lack of diagnostic/prognostic information and a distribution of features across the tissue biopsy that may lean towards one particular category may lead to the assignment of bridge ratings, which is consistent with some of the motivations for bridge category models.

### 4.1 Recommendations

From the simulation studies and real-world examples, we have a few guiding principles when selecting a DGM and appropriate corresponding model in the presence of bridge ratings:

1. In general, random stage assignments may likely contribute to biased effect estimates.
2. When the estimates acquired from up/down-staging differ, the mixture model may identify a mixture parameter estimate which can help explain the differences in effect. Relying on up or down-staging alone may be problematic when the true propensity for up/down-rating may be opposite to the ad-hoc approach.
3. If the effects from up/down staging are similar, the expanded model may provide a more precise estimate, but this is best accomplished in a situation where blurring effects are not suspected.
4. Reporting bridge ratings and interfacing with domain experts can contribute to an understanding of why they occur and inform approaches for proper adjustment. Should the domain experts communicate back that there was complete ambiguity in assignment, evaluating models under the expanded DGM may prove useful. The lack of mixing of the mixture parameter sampling may confirm this hypothesis. If the raters reply that instead, which was in our case for the real-world fibrosis data, that “stages assigned were mostly indicative of a 2, but had features of a 3”, then models may perform well under the assumption of the blurred DGM.

### 4.2 Limitations

These results, while promising, have limitations. We acknowledge that exhaustive tests over the simulations were not performed (asymmetric distributions of ordinal outcomes, different mass assigned to different uncertainty categories under the expanded DGM, interaction effects). However, the chosen simulations highlighted the advantages of the expanded and mixture approaches. Tests on the real-world data were conducted on two specific subsets of the data (two pathologists at particular testing intervals). While selection of the raters and tests were arbitrary, we did not model the nested (biopsies per patient) and cross-classified (repeat measures of biopsies versus pathologist) structure of the data in the real-world example in order to match the simplicity of the simulation analyses, which are focused more on methods development rather than application. In a true, real-world scenario, hierarchical Bayesian methods are employed to simultaneously adjust for multiple raters and repeated measurements [52]. We plan to extend the modeling approach into this context to account for rater specific effects and clustering by case and biopsy. In a similar vein, the mixture parameter utilized in the mixture model is a population parameter. While the effect explained by the parameter holds true across the cohort, the parameter in its current configuration is independent of any fixed or random effect and cannot currently use these factors to explain how they impact blurriness. For instance, blurriness patterns may be rater-specific, dependent on the assigned rating/conditional mean or be case-dependent (random intercept). Nor did we consider heteroskedastic variance [16], from which the variance in the response may be dependent on the covariates that estimate the conditional mean.

### 4.3 Opportunities

Opportunities exist to further develop and apply the method for consideration of more real-world scenarios, such as estimating variance parameters that describe between and within-cluster effects. Where appropriate, hierarchical modeling renditions of bridge category models may help assess the validity of the clinical grading/staging scale and the application in clinical trial design after proper validation [53].

While we have developed statistical models and estimation methods that are suitable to use with each data generating mechanism (DGM), we have not fully explored methods to selecting which model is best to use on a given data set for which the true DGM is unknown. We envision adapting Bayesian model comparison methods to this situation and to considering Bayesian model averaging methods; that latter will incorporate the uncertainty in the underlying DGM into the analysis. Such methods can make the posterior probability of each model being the true model a part of the output from the analysis. A yet further complication is that case when raters within a study may conform to different rating systems. While the observation of a bridge-rating implies what a rater may be thinking in terms of an expanded scale, the absence of one might imply that they are reporting on the original scale. However, one cannot identify whether the rating just happened to be a non-bridge rating or if the original scale was being used. Incorporating this uncertainty would extend the models into the latent class realm with the latent class being the choice of scale. Ideally this complication would be obviated at the design-stage by specifying to raters that bridge ratings should be used to represent truly discretionary assessments. However, this still does not offer absolute guarantee that the raters will exactly follow the stated protocol. Clearly, imposing some consistency across the use of a scale may be helpful. But as demonstrated by our results, if this resulted in raters arbitrarily determining their final ratings, this could potentially affect the quality of the data and so would not be a good practice.

Algorithms in artificial intelligence (AI) may resolve issues with interrater variability [54], measurement uncertainty and error by providing quantitative assessments of tissue histology and other tangential tasks which may be of immediate value [55, 12, 56, 57, 58, 59]. A combination of both rater uncertainty and incomplete information about the histology may present additional challenges for training and evaluating AI technologies. As such, these methods may benefit from incorporating such bridge category methods into their specification and evaluation. For instance, a machine learning model trained to stage a tumor may output an ordinal response. Such ordinal outcomes must be compared appropriately to a measurement from pathologist(s) which may include bridged stage assignments. As another example, a recent large scale validation study of virtual tissue staining technologies utilized down/upstaging as a strategy for overcoming bridged ratings [12]. Here, upstaging, downstaging, random-staging or treating the ordinal variable as continuous may violate the true DGM, leading to biased estimates and potentially over/understating the efficacy of the AI technology. Such studies may benefit from reassessment using bridge category modeling approaches. Finally, there exists opportunity to utilize such methods to reassess ordinal outcomes for molecular/omics data with bridge ratings [60, 61, 62, 24, 63]. As such, we plan to apply these methods towards better understanding their impact on clinical trial design, interrater variability for establishment of accepted grading/staging scales and development/assessment of AI technologies.

## 5 CONCLUSION

Ordinal ratings are commonly used in biomedical applications to assess factors related to disease progression. While such ratings are regularly employed for clinical decision making, inferring the effectiveness of grading/staging scales, evaluating clinical trial efficacy, and development and assessment of AI technologies, bridge ratings remain a relatively unexplored, ubiquitous phenomenon which may contribute to biased and imprecise study results if erroneously analyzed. While the data generating mechanisms may reflect scenarios where information is added/expanded, blurred, or collapsed, bridge ratings should be modeled, not dismissed. These ratings often have a specific meaning in terms of the precision or certainty with which a rater trusts their assessments (either higher or lower than the original scale may represent) and as such are a form of information. Failure to account for bridge ratings (reflecting greater uncertainty), implies a coarsening on the categories and a loss of information which will make the model work harder than necessary to fit that observation. In turn, this will reduce the relative attention the model applies to other observations whose precision is greater and again will result in a lack of efficiency compared to the optimal analysis. Under both scenarios, ad hoc methods that truncate, randomly re-assign, or otherwise transform the data to get it into a form amenable to a standard analysis may impart bias on the results. By building and using statistical models that account for the believed data generating mechanism(s), we can potentially improve practices surrounding drug approval, validation of measurement scales and evaluation of diagnostic decision aids. At a minimum, reporting bridged ordinal ratings and discussing with domain experts and raters as to how such ratings arise should be actively encouraged in biomedical research.

## acknowledgements

We would like to acknowledge Christian Haudenschild, Julian Gullett, Todd MacKenzie and Jorge Lima for their thoughtful discussion on the subject matter.

This work was supported by:

- NIH grants R01CA216265, R01DE022772, and P20GM104416 to BCC.
- Two Dartmouth College Neukom Institute for Computational Science CompX awards to BCC, LJV and JJL.
- JJL and CB are supported through the Burroughs Wellcome Fund Big Data in the Life Sciences training grant at Dartmouth.
- Norris Cotton Cancer Center, DPLM Clinical Genomics and Advanced Technologies EDIT program.
- NIH grant P01AG019783 to support AJO.

## Bridged Category Model Appendix

### 1 ORDINAL RESPONSE VARIABLES AND CUMULATIVE LINK MODELS (CLM)

An ordinal response variable may reflect a latent continuous process, the categories of which are represented by adjacent intervals of differing width within the distribution. While it is erroneous to directly encode ordinal variables as nominal or continuous [1, 2, 3, 4, 5], several regression techniques have been developed to represent ordinal data[6]. The cumulative link model is a special case of ordinal regression models and is particularly advantageous in that it bears resemblance to both categorical and continuous processes through indirect modeling of a latent continuous process (Main Text Figure 1A).

Let *Y* represent the ordinal response variable, *j* ∈ {1, 2,. . ., *K* - 1} (i.e., disease stage). Let *X* be a design matrix of observations by predictors. The latent variable, *Y*\*, represents the true, unobserved continuous process (i.e., disease progression) underlying the ordinal observations. The predictors serve to explain this latent variable. Thus, a linear combination of these predictors may be employed to generate:

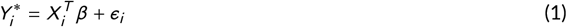

where *ϵ*_*i*_ ~ *N* (0, 1) is the parametric distribution assumption consistent with the ordered probit regression model. Alternatively, the logistic link function is obtained by exchanging the assumption of normality of the error terms for the logistic distribution, yielding the proportional odds model. Generally, the error term may have any cumulative distribution *F*, which impacts model fitting and interpretation of the results. The conditional mean is given by:

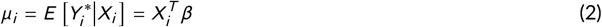

Cumulative link models (CLM) include a set of threshold parameters, {*θ*_*j*_}_*j*∈{1,2,...,*K*-1}_, which correspond to cutpoints of a continuous latent distribution from which partially observed or limited outcomes are obtained. These also define endpoints from which interval and continuous probasbilities are defined, from which the effect of covariates can be measured against. Intervals between the cutpoints represent the observed classes {*Y*_*j*_}_*j*∈{1,2,...,*K*}_. Here, we include the motivation and derivation of the link function and likelihood. The thresholds, represented by vector 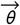 are used to partition the latent distribution as:

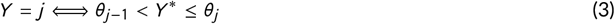

In this latent distribution, distances are meaningfully encoded, which permits the use of linear regression on the latent outcome:

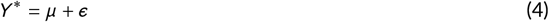

where *μ* represents the conditional mean, and *ϵ* the error term as above. In order to map the latent distribution to a set of probabilities, we specify the distribution *F* of *ϵ* (which can be any cumulative distribution):

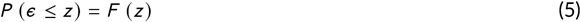

Combining the previous three equations:

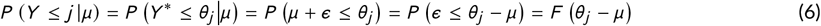

The probability of *Y* being less than or equal to *j* given the conditional mean *μ* is equal to the cumulative distribution *F* evaluated at *θ*_*j*_ - *μ*. Because the probability of an event occuring and the probability of it not occurring sum to 1, it follows that:

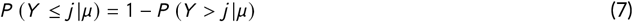

Therefore, the probability *Y* = *j* given *μ* is as follows:

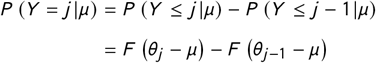

Finally, the likelihood of an observation *Y* = *j* assuming it is generated from a multinomial distribution with a cumulative link function defining category probabilities as above is given by:

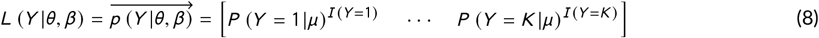

We summarize the likelihood and link function below:

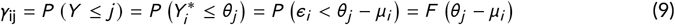

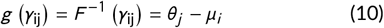

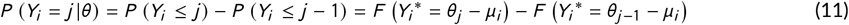

In this model, the probability of drawing a particular class is the area of the cumulative distribution between two estimated cutpoints after centering by *μ*_*i*_. At the tails of the distribution, where *Y* ∈ {1, *K*}, the predicted probabilities of the classes are *P*(*Y* = 1) = *F*(*Y*_*i*_* = *θ*_1_ - *μ*_*i*_) and *P*(*Y* = *K*) = 1 - *F*(*Y*_*i*_* = *θ*_*K*-1_ - *μ*_*i*_) respectively. The threshold and conditional mean parameters of these models are not identifiable, as the conditional mean *μ*_*i*_ can be re-scaled without changing the fitted probabilities, given the constraint of the cumulative distribution *F* (i.e. multiplying *θ* - *μ* by a constant leads to an equivalent fit) [7]. However, by imposing the restriction that the observations have a variance of 1 (or any other constant), we can make the model parameters identifiable by the data. We implemented estimation of a multinomial model with the above cumulative link function using the Stan probabilistic programming language [8, 9]. Stan uses Hamiltonian Monte Carlo methods to perform Bayesian model estimation (See Appendix, section “Bayesian Computation and Hamiltonian Monte Carlo”) [10, 11]. The Bayesian model for a cumulative link model for *K* ordinal response categories is specified below for predictors indexed by *m* ∈ {1, 2,. . ., *M*} and cutpoints from *j* ∈ {1, 2, 3,. . ., *K* - 1}:

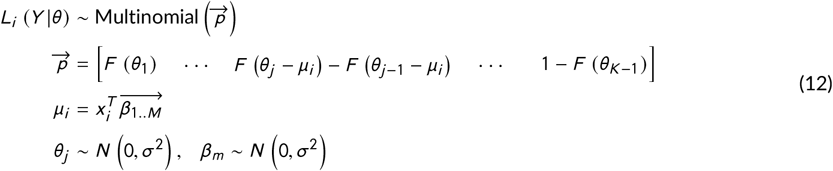

### 2 BAYESIAN COMPUTATION AND HAMILTONIAN MONTE CARLO

While frequentist statistics focus on optimization of the likelihood, Bayesian statistics focus on estimation of the posterior distribution, *p*(*θ*|*X*), where *θ* are parameters treated as random variables and *X* are the data that the posterior is conditioned on and thereby are treated as fixed [12, 5]. The posterior can be related to the likelihood of the cumulative link model via Bayes Theorem:

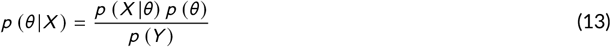

In eq.(13) *p*(*θ*) is the prior distribution of the parameters and *p*(*Y*) is the marginal distribution of the data, a normalizing factor. The Bayesian approaches were essential for estimation of our models because of their flexibility to handle different DGM, specification of priors, and better estimates of uncertainty through sampling the posterior rather than some approximation through maximum likelihood methods. Bayesian priors centered around zero may potentially also make estimates more conservative, which can in turn reduce the need for correction for multiple comparisons.

The cumulative link model, as implemented in packages, such as <u>clm</u> [13], typically involves optimization of the likelihood, given some starting parameters. Here, we utilize Bayesian model fitting procedures versus the frequentist approaches because estimates avoid approximations to aid computation, are typically more precise, numerical computations may be easily implemented and there is no strict constraint on the shape of the posterior distribution of the unknown parameters. While frequentist approaches optimize the likelihood for a set of parameters using maximum likelihood estimation and approximate the variance of the estimator using the Hessian, Bayesian approaches derive a posterior distribution for a set of parameters by updating the prior hypothesis with appropriate weight given to the likelihood over the prior. Amongst a few different approaches towards posterior estimation (eg. variational inference), Markov Chain Monte Carlo (MCMC) methods sample the posterior distribution of parameters until convergence of the joint posterior to a stationary solution. Hamiltonian Monte Carlo, while not conditionally sampling posterior parameters via a Markov Chain, is similar in spirit as the MCMC algorithms, drawing from the joint density of the model parameters and some auxiliary momentum variables, sampling through simulation of moving particles for a number of time steps under the joint state (*θ*) and momentum (*ϕ*) density: *p*(*ϕ*, *θ*) = *p*(*ϕ*|*θ*) *p*(*θ*), where the goal is to ultimately sample the joint posterior *p* (*θ*). In our motivating example (e.g., CLM), 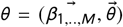 and 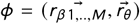, where *r* corresponds to parameter-specific momentum parameters, initially drawn from a uniform distribution then updated over time by the Hamiltonian dynamics of the system. The Hamiltonian is represented by:

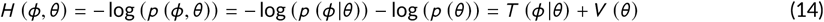

Where T and V represent the kinetic and potential energy respectively. Hamilton’s equations govern how particles move across the density landscape, which can be summarized as:

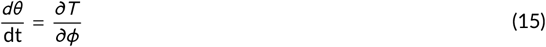

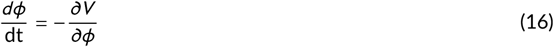

Automatic numerical differentiation solves the above differential equations, allowing the particles to move for <u>L</u> steps given a random sampling of momentum to initialize the trajectory, where they are then sampled (collection of parameters *θ*) and either accepted or rejected based on Metropolis Hastings criteria. No U-Turn Sampling (NUTS) is an extension on this sampler that attempts to increase the efficiency of the sampling process through adaptive measures that promote landscape exploration. We have implemented such samplers using the Stan programming language, integrated into R via <u>RStan</u> [9, 10], with some elements of the code inspired by design choices featured in the <u>brms</u> package [8].

### 3 LOCATION OF EXPANDED SUPPLEMENTARY TABLES

Expanded supplementary tables detailing assessment of model performance may be found in additional file 1. The first tab provides descriptions of the sensitivity tests for the simulation studies, the data generating mechanisms and the true covariate effects.

### 4 SUPPLEMENTARY RESULTS TABLES AND FIGURES

**APPENDIX TABLE 1.** Tabulated stage assignments for pathologists at particular tests

| Pathologist | 0 | 0-1 | 1 | 1-2 | 2 | 2-3 | 3 | 3-4 | 4 |
| --- | --- | --- | --- | --- | --- | --- | --- | --- | --- |
| Pathologist 1 | 10 | 12 | 33 | 10 | 52 | 31 | 56 | 24 | 28 |
| Pathologist 2 | 2 | 8 | 18 | 10 | 82 | 26 | 58 | 19 | 33 |

**APPENDIX TABLE 2.** Posterior estimates for mixture parameters for fibrosis staging model (BMI, Fib4, AST:ALT Ratio) for different pathologists

| Pathologist | $\bar{p}$ | 90% CI Low | 90% CI High | Posterior Interval Width |
| --- | --- | --- | --- | --- |
| 1 | 0.91 | 0.665 | 1 | 0.335 |
| 2 | 0.148 | 0 | 0.472 | 0.472 |

**APPENDIX FIGURE 1.**
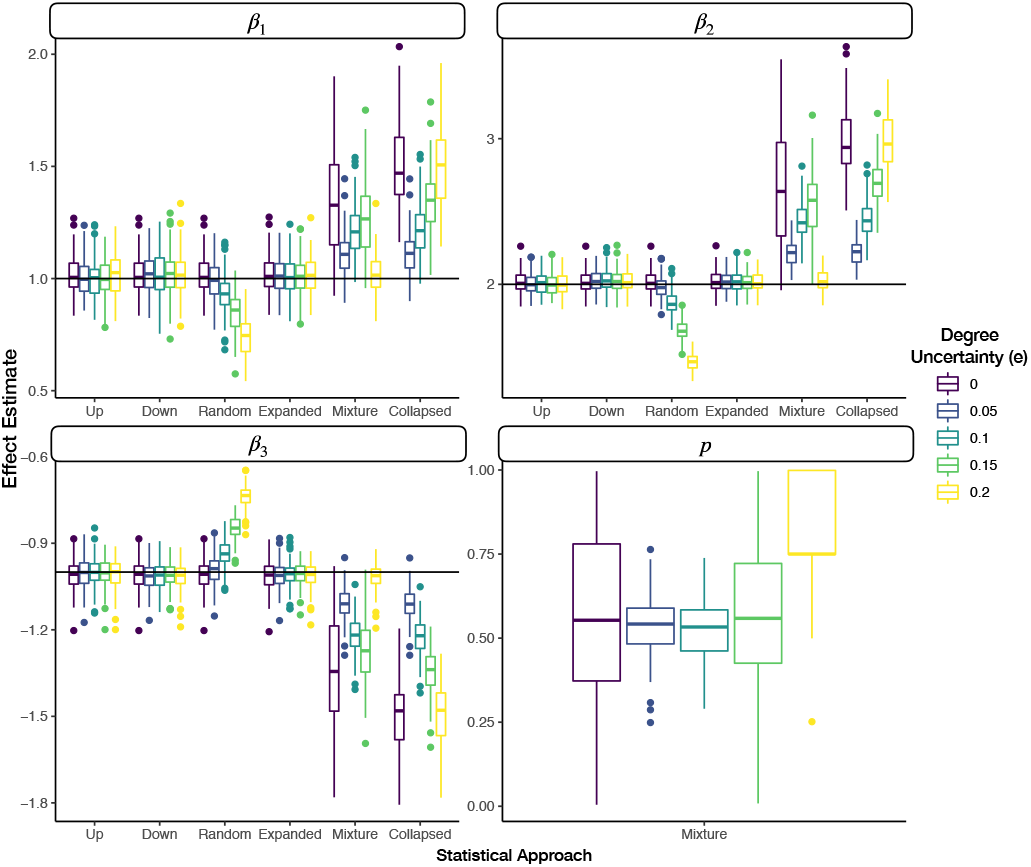
Faceted grouped boxplots indicating distribution of mean posterior covariate and mixture parameter effect estimates across simulated datasets; each facet presents different covariate/mixture effect; x-axis labeled by which data augmentation/modeling approach was fit to the data; y-axis is effect estimate; horizontal line added to indicate true population parameter for each covariate; boxplots are colored by the degree of uncertainty in the expanded DGM, where the effect size of the binary covariate is 1; all covariates were standardized prior to generation of the ratings

**APPENDIX FIGURE 2.**
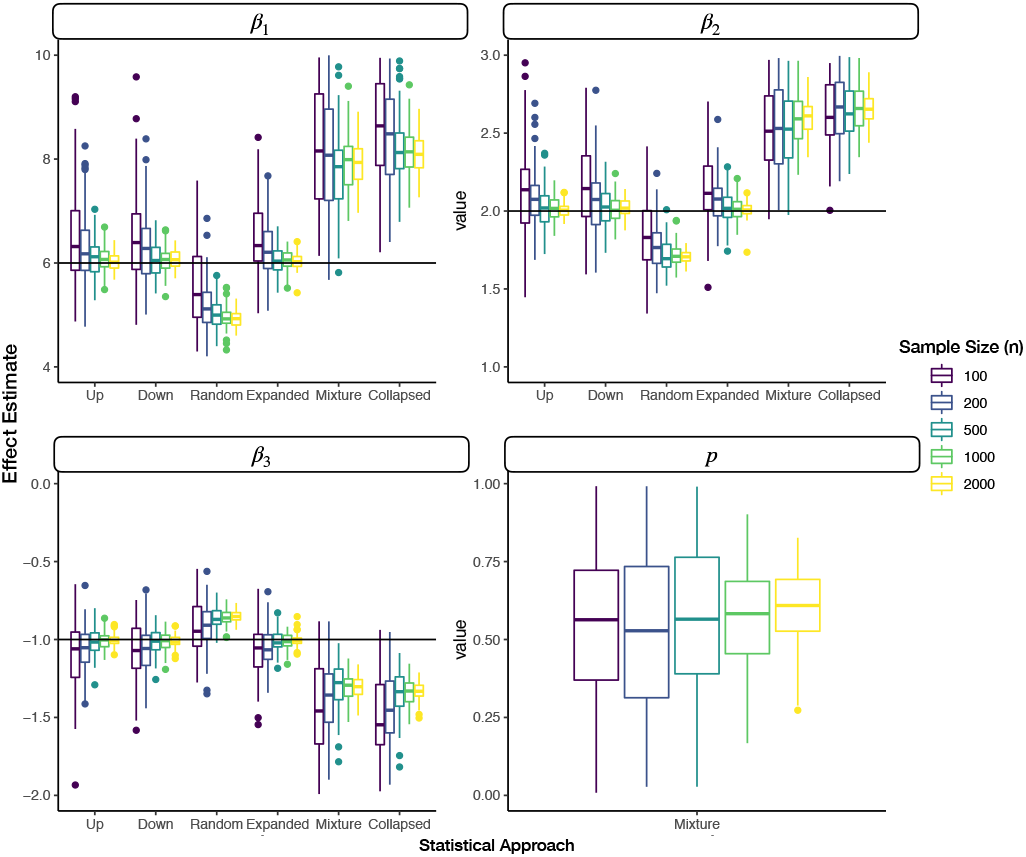
Faceted grouped boxplots indicating distribution of mean posterior covariate and mixture parameter effect estimates across simulated datasets; each facet presents different covariate/mixture effect; x-axis labeled by which data augmentation/modeling approach was fit to the data; y-axis is effect estimate; horizontal line added to indicate true population parameter for each covariate; boxplots are colored by the sample size in the expanded DGM, where the effect size of the binary covariate is 6

**APPENDIX FIGURE 3.**
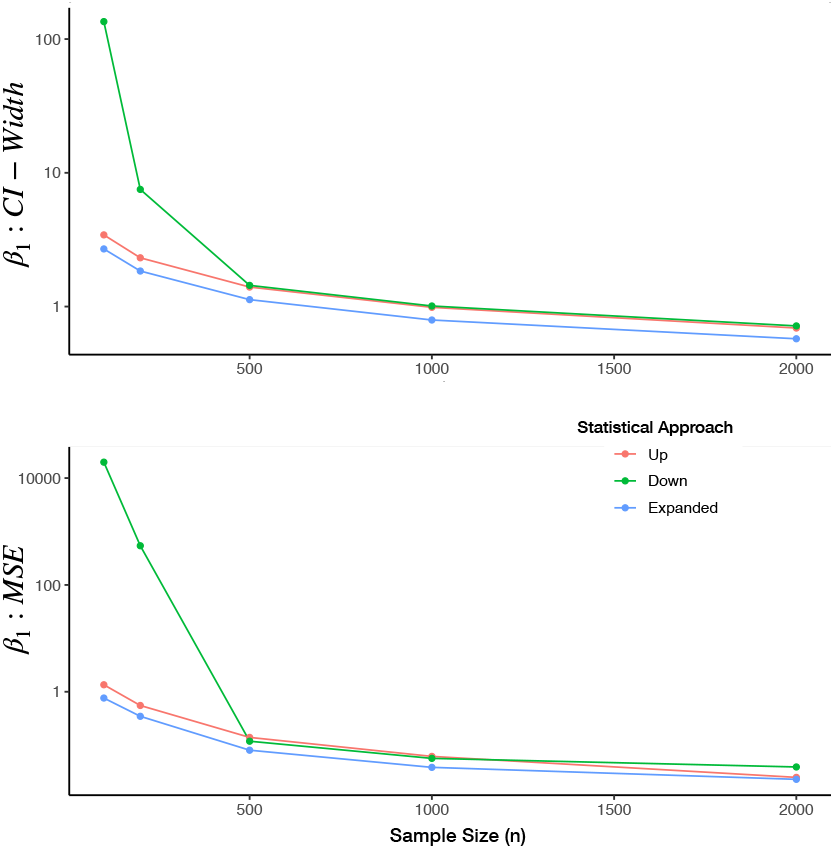
Sample Size (n) versus posterior interval width and MSE for the up, down and expanded category models for true effect size of six for the binary covariate

**APPENDIX FIGURE 4.**
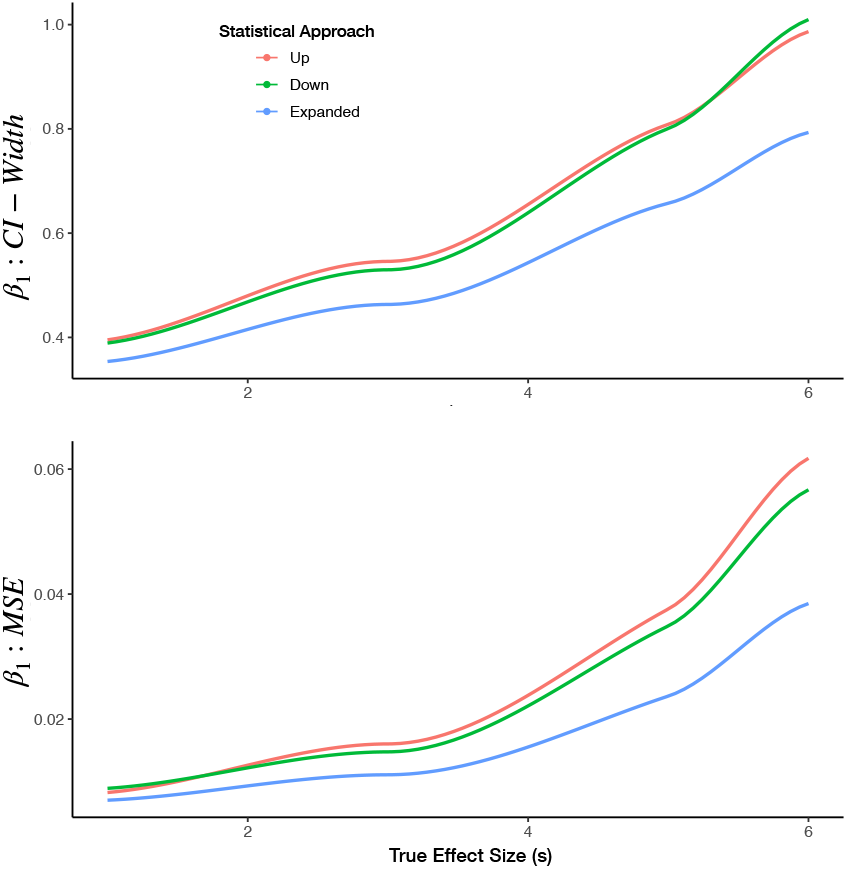
True Effect Size (s) versus posterior interval width and MSE for the up, down and expanded category models; results reported for the binary covariate and smoothed using loess regression

**APPENDIX FIGURE 5.**
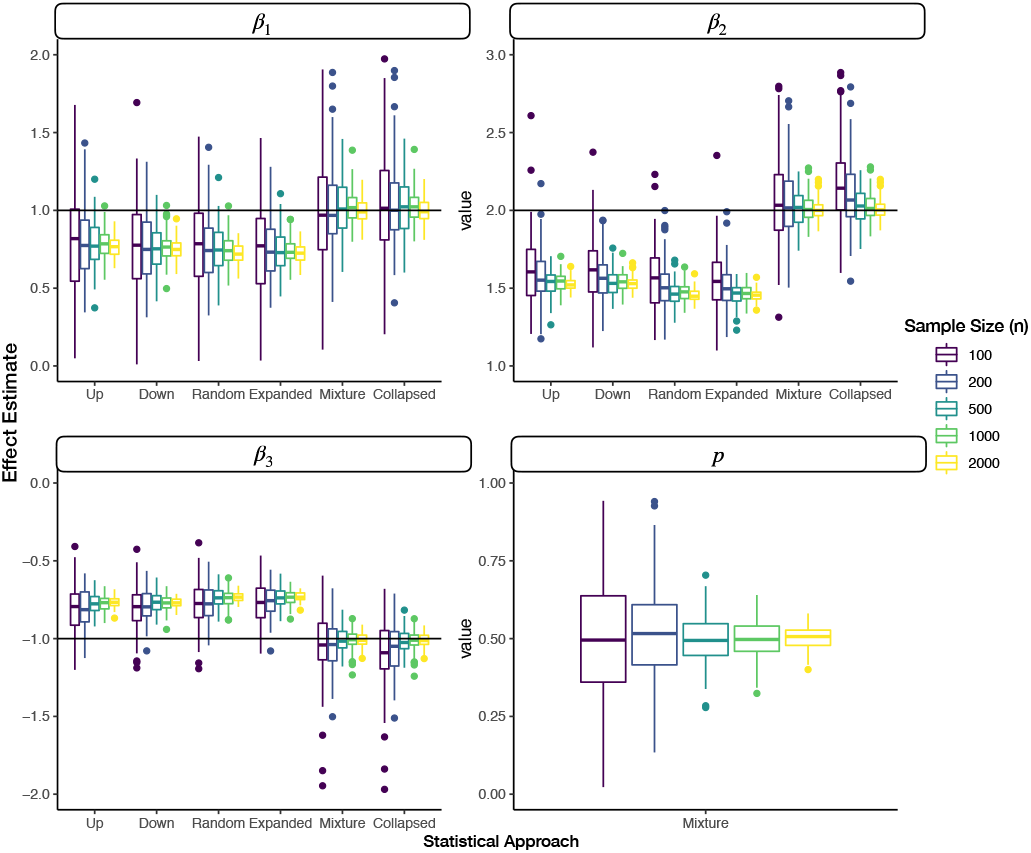
Faceted grouped boxplots indicating distribution of mean posterior covariate and mixture parameter effect (true mixture effect of 0.5) estimates across simulated datasets; each facet presents different covariate/mixture effect; x-axis labeled by which data augmentation/modeling approach was fit to the data; y-axis is effect estimate; horizontal line added to indicate true population parameter for each covariate; boxplots are colored by the number of samples for the blurred DGM

**APPENDIX FIGURE 6.**
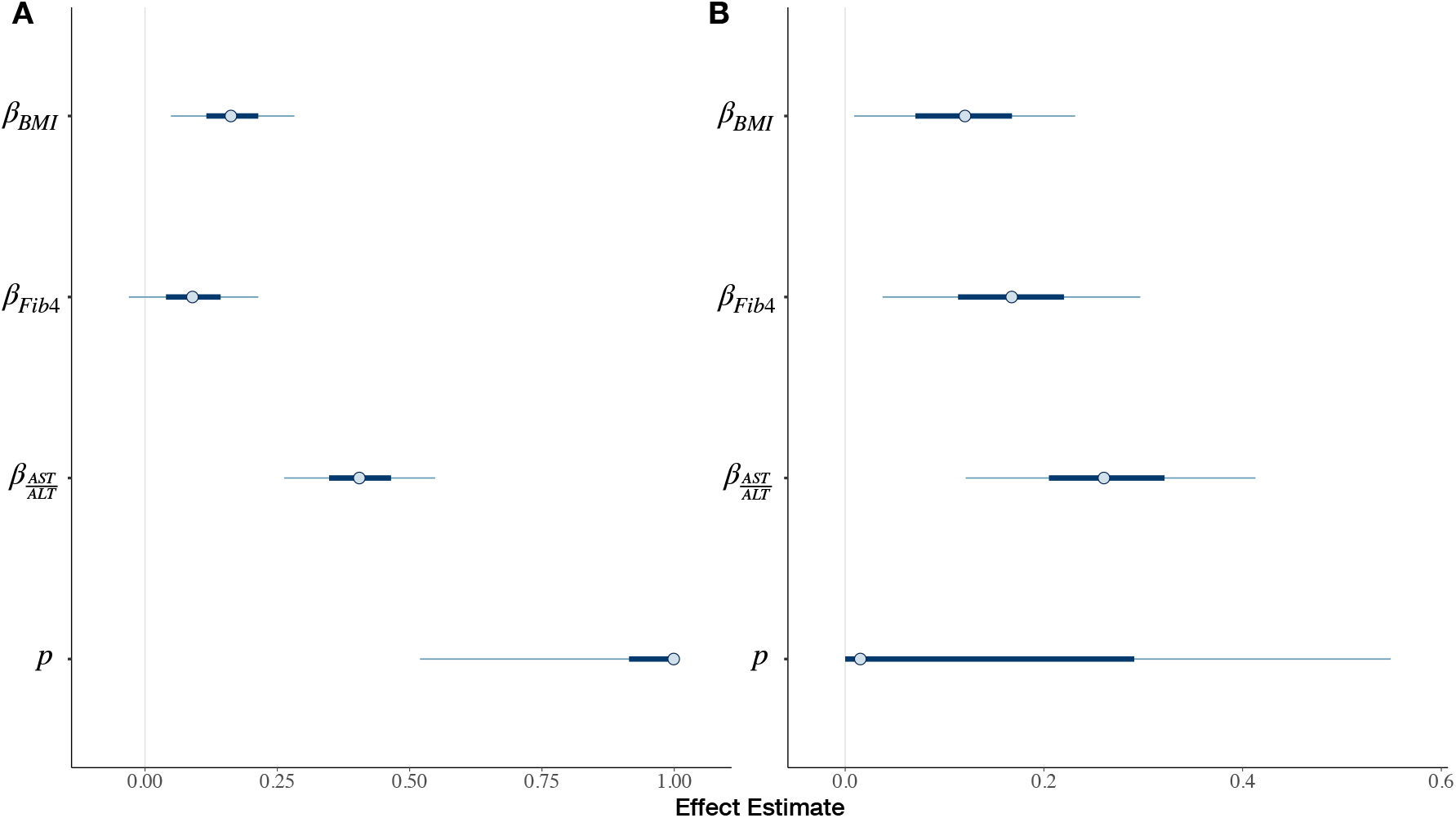
Plot of posterior intervals of covariates (BMI, Fib4, AST:ALT ratio) and mixture parameters for <u>Mixture Adjacent Model</u> for: a) Pathologist 1, and b) Pathologist 2

### 5 ADDITIONAL DISCUSSION

#### 5.1 Expanded DGM Simulation

For the expanded DGM, the expanded approach provided greater precision in estimates versus up and down staging. This was true across all sample sizes and effect sizes. However, greater precision is conferred when the sample size is small, effect size is large, and the proportion of ratings assigned to a bridge rating is similar to that of the certain assignments. We were surprised to find that ordinal models fit on up- and down-staged responses to exhibit similar degrees of bias as the expanded approach. However, these results are consistent with prior studies which demonstrate that collapsing adjacent categories has negligible impact on bias but inflates the standard error of the estimate [14, 15, 16, 17, 18]. Likewise, the expanded categories are able to maintain high precision, while down and up rating provides coarser estimates of the true ratings. Random up/down rating may serve to bias and deflate the effect estimate toward zero, while the mixture and collapsed approaches may inflate the effect estimate. As such, when confronted with data that is suspected to be of the expanded DGM, the expanded model offers the truest estimate of the effect.

#### 5.2 Blurred DGM Simulation

For the blurred DGM, we note vastly superior performance for the mixture and collapsed models as compared to the traditional approaches, regardless of the degree by which the ordinal ratings were blurred, the propensity for assigning the upper two categories, sample size and effect size. The mixture model would appear to learn the properties of the data that it is fitting, while the collapsed category was highly applicable when the propensity of bridge-ratings is around 0.5, but slightly biased otherwise. This is likely because the collapsed category model considers two categories simultaneously, much like the mixture model but does not learn a mixture parameter. The flexibility in learning a population parameter avoids bias when measurements are blurred. In contrast, the effect estimates provided by the expanded and traditional approaches were biased towards zero. As such, when confronted with data that is suspected to be of the blurred DGM, the mixture adjacent model offers the truest and most flexible estimate of the effect.

#### 5.3 Deciding on the Ideal DGM

In a real-world scenario, it can be difficult to decide which DGM is applicable to the data. Likelihood ratio tests calculated on the data for model selection may lead to misleading conclusions when the ratings are coarsely treated [19, 20, 21]. Constructing a likelihood test (e.g. Bayes Factor and Pareto Smoothed Importance Sampling) to compare the expanded model to the other approaches is challenging due to the fact that one model is not nested in the other and the two models support non-overlapping categories. In sum, a mixture of interviewing expert raters and data driven testing of parameters and comparison of model effects to differences in effects registered under simulations should inform the selection of both the DGM and ideal model.

#### 5.4 Additional Limitations and Opportunities

While our models took into account bridged categories, they do not explicitly account for instances of greater measurement error (outside of those expected from adjacent categories). However, given the widely varying reports of stages between the two raters (high inter-rater variability), this is suggestive of additional measurement error which may have obfuscated some of the effects of the true DGM [22]. Potentially, bridge category models may benefit from incorporating elements from other models which tackle uncertainty across larger number of categories. For instance, mixture adjacent models bear resemblance to the proposed <u>GEM</u> (Generalized Mixture Models with Uncertainty) and <u>CUB</u> (Combination of a discrete Uniform and a shifted Binomial random variable), models and their derivatives [23, 24, 25]. These models attempt to ascribe two components to providing ordinal ratings. Feeling (attraction and awareness) components towards a particular rating may make the rating more likely to be chosen, as modeled by a Binomial random variable. Uncertainty (indecision and blurriness) components, from which a discrete uniform distribution is assumed, places greater weight on the remaining categories. However, neither of these methods take into account bridge ratings into the decision-making process and are thus inappropriate for the treatment of bridge ratings yet feeling and uncertainty components may inform future iterations of bridge category models.

In addition, the DGM may be rater-specific, as we had estimated different mixture parameters for different pathologists, which may be correspondent to the widely varying effect estimates within and between raters depending on the model being used to fit the data. Given the relatively moderate degree of bridged category assignment, these were not of high enough magnitude for effects to become heavily distorted. While the effect of inter-rater variability more likely pertains to measurement error and diminishing of effect estimates, utilizing the mixture approach in one instance in the real-world setting provided a significant effect estimate, when the other approaches had suggested otherwise.

